# Varying-Censoring Aware Matrix Factorization for Single Cell RNA-Sequencing

**DOI:** 10.1101/166736

**Authors:** F. William Townes, Stephanie C. Hicks, Martin J. Aryee, Rafael A. Irizarry

**Affiliations:** Department of Biostatistics, Harvard T.H. Chan School of Public Health, Boston, MA; Department of Biostatistics and Computational Biology, Dana-Farber Cancer Institute, Boston, MA; Molecular Pathology Unit, Massachusetts General Hospital, Charlestown, MA; Center for Cancer Research, Massachusetts General Hospital, Charlestown, MA; Department of Pathology, Harvard Medical School, Boston, MA

## Abstract

Single cell RNA-Seq (scRNA-Seq) has become the most widely used high-throughput technology for gene expression profiling of individual cells. The potential of being able to measure cell-to-cell variability at a high-dimensional genomic scale opens numerous new lines of investigation in basic and clinical research. For example, by identifying groups of cells with expression profiles unlike those observed in cells with known phenotypes, new cell types may be discovered. Dimension reduction followed by unsupervised clustering are the quantitative approaches typically used to facilitate such discoveries. However, a challenge for this approach is that most scRNA-Seq datasets are sparse, with the percentages of measurements reported as zero ranging from 35% to 99% across cells, and these zeros are partially explained by experimental inefficiencies that lead to censored data. Furthermore, the observed across-cell differences in the percentages of zeros are partly due to technical artifacts rather than biological differences. Unfortunately, standard dimension reduction approaches treat these censored values as true zeros, which leads to the identification of distorted low-dimensional factors. When these factors are used for clustering, the distortion leads to incorrect identification of biological groups. Here, we propose an approach that accounts for cell-specific censoring with a varying-censoring aware matrix factorization (VAMF) model that permits the identification of factors in the presence of the above described systematic bias. We demonstrate the advantages of our approach on published scRNA-Seq data and confirm these on simulated data.

## 1 Introduction

High-throughput technologies provide transcription measurements for each gene, resulting in tens of thousands of measurements for each biological sample. Currently, RNA sequencing (RNA-Seq) is the most widely used of these technologies. Due to the experimental protocol’s requirements, to obtain reliable measurements one must provide hundreds of thousands to millions of cells as input. The resulting measurements are therefore averaged expression levels across the many cells that form the biological specimen of interest. In contrast, single cell RNA-seq (scRNA-Seq) protocols can produce measurements with material from just one cell as input. To clearly distinguish these two technologies we refer to the former as bulk RNA-Seq.

A promising application of scRNA-Seq technology is the identification of sub-populations of cells within biological samples. This is particularly interesting due to the possibility of discovering previously unknown cell types or differences between cells previously thought to be indistinguishable. For example, previous studies have sought to identify new cell types or transcriptional heterogeneity in tissues such as brain [Zeisel et al., 2015], blood [Villani et al., 2017], melanoma [Tirosh et al., 2016], glioblastoma [Patel et al., 2014], retina [Macosko et al., 2015], and spleen [Jaitin et al., 2014].

Consider a scRNA-Seq experiment that measures transcription profiles in hundreds to thousands of individual cells. We can represent each of the *N* cells as a *G-*dimensional vector with *G* as the number of genes. An unsupervised clustering analysis is often applied to organize cells into groups based on the similarity of their expression profiles. Because *G* is in the thousands, the clustering is typically aided by first reducing the dimensionality. Dimension reduction techniques commonly applied in published scRNA-Seq studies include linear methods such as Principal Components Analysis (PCA) and nonlinear methods such as t-distributed Stochastic Neighbor Embedding (t-SNE), [van der Maaten and Hinton, 2008]. These techniques rely on defining a distance metric to quantify pairwise similarities between cells.

A challenge in applying dimension reduction methods to scRNA-Seq data is that the proportion of genes reported to be exactly zero in scRNA-Seq data tends to be substantially higher than in bulk RNA-Seq datasets. The larger proportions of zeros make methods motivated by Gaussian models inappropriate. In particular, because data is log transformed after adding a constant to avoid logging zeros, the choice of this constant, which will typically be the mode of the data distribution, will greatly affect the estimation of factors. To account for the larger number of zeros, Pierson and Yau [2015] developed Zero Inflated Factor Analysis (ZIFA). This approach models unobserved expression levels with factors and assumes that the observed data includes zeros due to censoring. However, the censoring model includes only a single parameter shared across all cells and therefore, the model does not explicitly address the systematic errors that result in different censoring behavior across different cells.

Hicks et al. [2017] performed a comprehensive re-analysis of published data and found that the proportions of zeros ranged from 35% to 99% across cells and that in a substantial number of studies, the proportion of zeros was highly correlated with the first principal component of the gene expression data. Furthermore, Hicks et al. [2017] provide evidence that differences in the proportion of zeros is frequently driven by technical artifacts rather than biological factors. We demonstrate that failing to address these features of the data can lead to identification of spurious clusters even when the underlying data are essentially homogeneous. Here, we propose an approach we refer to as Varying-Censoring Aware Matrix Factorization (VAMF), which models cell-specific detection patterns using random effects. We demonstrate the advantages of VAMF over current approaches with published datasets as well as with simulations.

To motivate our approach, we downloaded three datasets that included paired bulk and single cell measurements on the same biological samples, namely those used in Shalek et al. [2013], Trapnell et al. [2014], and Patel et al. [2014]. The bulk data served as an independent measurement used to explore the relationship between the censoring mechanism and the underlying expression levels.

One of these datasets, Patel et al. [2014], examined five distinct biological specimens, specifically five glioblastoma tumors from five different individuals, and included a subset in which cells taken from the same biological specimen were processed in two different batches. We refer to this dataset as the *glioblastoma* dataset and to the subset as the *two-batches* dataset.

We represent the normalized scRNA-Seq data with *Y_ng_*, with *n* = 1, …, *N* indexing the cells and *g* = 1, … *,G* indexing the genes. We denote the normalized bulk RNA-Seq data with *M_g_*. We define 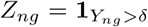 to be a binary detection indicator. Detection occurs if the observed expression is above the threshold *δ* ≥ 0. Following Shalek et al. [2013] and Hicks et al. [2017] we set *δ* = 1 normalized count, which accounts for low magnitude Poisson noise [Kharchenko et al., 2014] also referred to as ‘shot noise’ [Matz et al., 2013]. We define 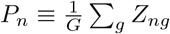 or the fraction of genes in cell *n* that were detected above the threshold and refer to this quantity as the *detection rate*. We define a similar quantity for the bulk RNA-Seq. Specifically we define 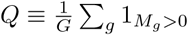.

A widely used method for class discovery is to apply PCA or t-SNE to transformed data log_2_(*Y*_*ng*_ + 1). Using these approaches on the *two-batches* dataset, we see two clear clusters (Figures 1A-C). Therefore, if we were following the current state-of-the-art analysis this would lead us to conclude that there are at least two distinct cell types in this biological specimen. Unfortunately, the two clusters correlate strongly with the batches, which leads us to conclude that this is a false discovery. Furthermore, the distributions of *P_n_* differ strongly across technical batch (Figure 2A) and correlate strongly with the leading principal components (Figures 2B-C). This suggests that the proportion of zeros, which differ substantially across the batches, are driving these results.

**Figure 1:**
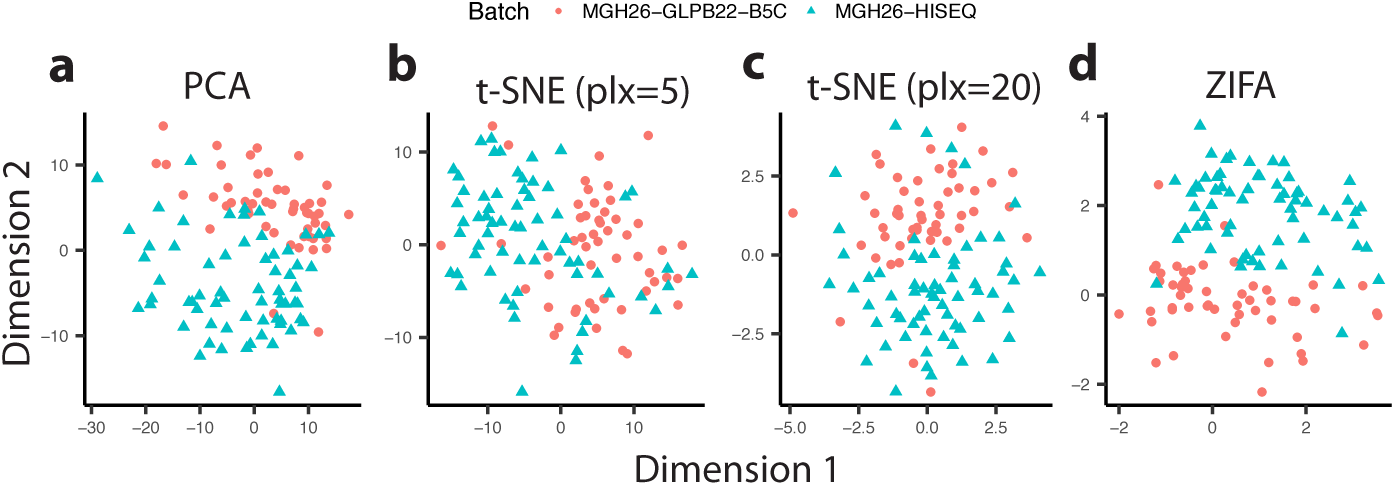
Dimension reduction results for the *two-batches* dataset. The two batches are represented by two colors. We show results for four methods: A) Principal Components Analysis (PCA), B) t-distributed Stochastic Neighbor Embedding (t-SNE) with perplexity (plx) parameter 5, C) t-SNE with perplexity parameter 20, and D) Zero Inflated Factor Analysis (ZIFA).

**Figure 2:**
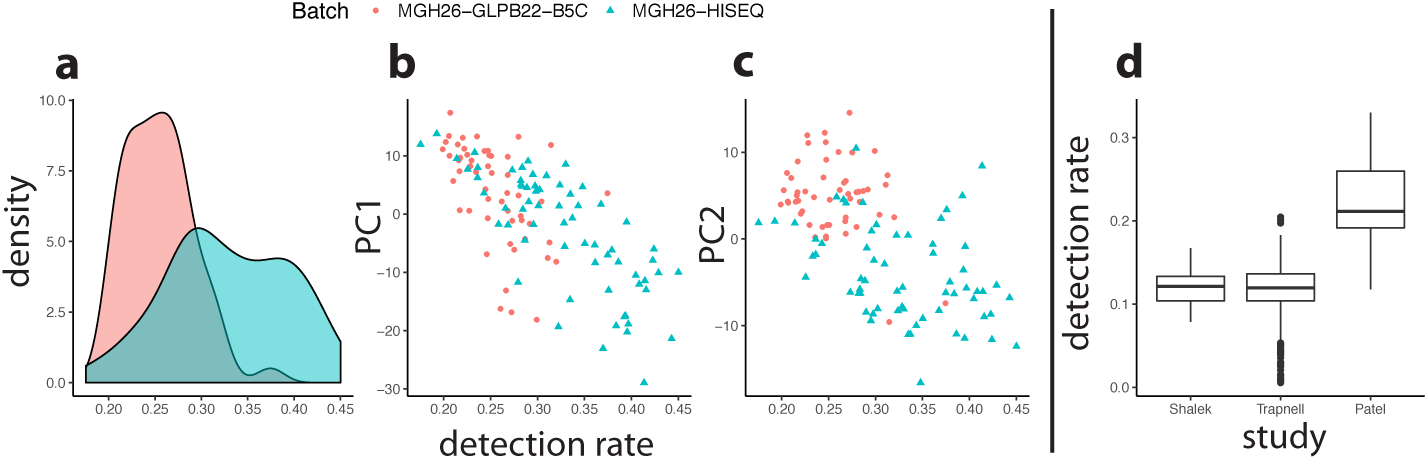
Detection rates drive distance calculations. A) Smooth densities for the detection rates in the *two-batches* datasets with color representing batch. B) First principal component versus detection rate. C) Second principal component versus detection rate. D) Distributions of single cell detection rates stratified by three scRNA-Seq studies.

Using ZIFA leads to the same incorrect conclusion (Figure 1D), therefore modeling the zeros as censored is not enough to avoid the false discovery. ZIFA accounts for censoring using the model:

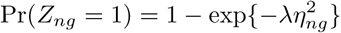

where *η_ng_* is the unobserved true log-expression value and λ controls the severity of censoring. For the non-censored observations, the ZIFA model is:

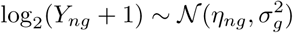

Finally, a standard factor analysis model is placed on the latent expression 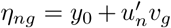 where *u_n_, v_g_* ∈ ℝ^*L*^. By using only one parameter for censoring, λ, the ZIFA model is unable to capture the variation in detection rate *P_n_* across cells and this variation appears to bleed through into the factors (Figure 1D).

Note that detection rate variability is common in all experiments that we assessed, with *P_n_* typically ranging from 0.1 to 0.4 (Figure 2D). Also note that *Q* is .28 for Trapnell et al. [2014], .36 for Shalek et al. [2013], and .6 for the *two-batches* dataset [Patel et al., 2014]. Therefore, the detection rate in scRNA-Seq is much smaller than in bulk RNA-Seq.

The relationship between *P_n_* ≪ *Q, n* = 1,…, *N* is expected since a gene that is considered to be expressed in a tissue need not be expressed in every single cell at all times for *M_g_* > 0. However, Hicks et al. [2017] demonstrated that part of the *P_n_* variability is explained by technical reasons. It is therefore indispensable that we model this varying censoring mechanisms appropriately.

## 2 Results

### 2.1 Joint Likelihood Model for Censored and Detected Data

We denote the unobserved log-transformed expression data with *η_ng_* and similar to ZIFA we model it using latent factors:

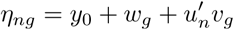

Here *y*_0_ represents a global intercept and *w_g_* represents a gene or feature-specific effect. We do not include a sample-specific global effect since these should be accounted for by normalization. The latent sample and feature factors are represented with *u_n_*, *v_g_* ∈ ℝ^*L*^ with *L* ≪ min{*N, G*}. The sample factors *u_n_*, are the key target of inference since these may be used to group cells. The *u_n_* are analogous to principal components if not for the censored data. The *v_g_* are analogous to loadings in PCA.

The key part of our model is the censoring mechanism *f_n_*(*η_ng_*) ≡ Pr(*Z_ng_* = 1 | *η_ng_*). We used the bulk data to motivate a parametric model for *f_n_*(*η_ng_*). Although due to censoring *η_ng_* was unobservable, we could observe *M_g_* and expected that E[log(*M_g_*)] = E[*η_ng_*] since the bulk tissue contained millions of cells from the same specimen. We therefore assumed *f*_*n*_(*η*_*ng*_) ≈ Pr(*Z_ng_* | *M_g_*). According to Bengtsson et al. [2008] and Reiter et al. [2011], the reverse transcription process (an essential precursor to RNA-Seq) is sensitive to low levels of starting material (RNA) in individual cells. Intuitively, if too few RNA copies are present, the chance of zero transcripts being measured (censoring) increases greatly. This suggested *f_n_* to be non-decreasing.

Nonparametric estimates (Figure 3) of *f*_*n*_ were obtained using non-decreasing splines comparing *Z_ng_* versus log_2_(*M_g_*) for *M_g_* > 0. Across all three experiments, the same sigmoid pattern was observed, motivating the use of logistic curves to model *f*_*n*_. Furthermore, the spline fits implied that the slopes and intercepts should be allowed to vary from cell to cell. We therefore use the following model for the censoring mechanism:

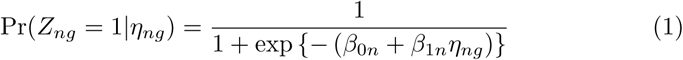

The parameters *β*_0*n*_ and *β*_1*n*_ account for the fact that each cell has a different censoring mechanism. This feature makes our model fundamentally different from ZIFA. Observed data *Y_ng_* are modeled using the likelihood:

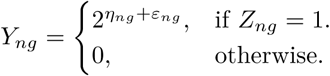

with *ε*_*ng*_ independent and identically normally distributed random variables with mean 0 and standard deviation *σ_y_*. The log-normal distribution of gene expression has been justified by McDavid et al. [2013] as matching empirical distributions.

**Figure 3:**
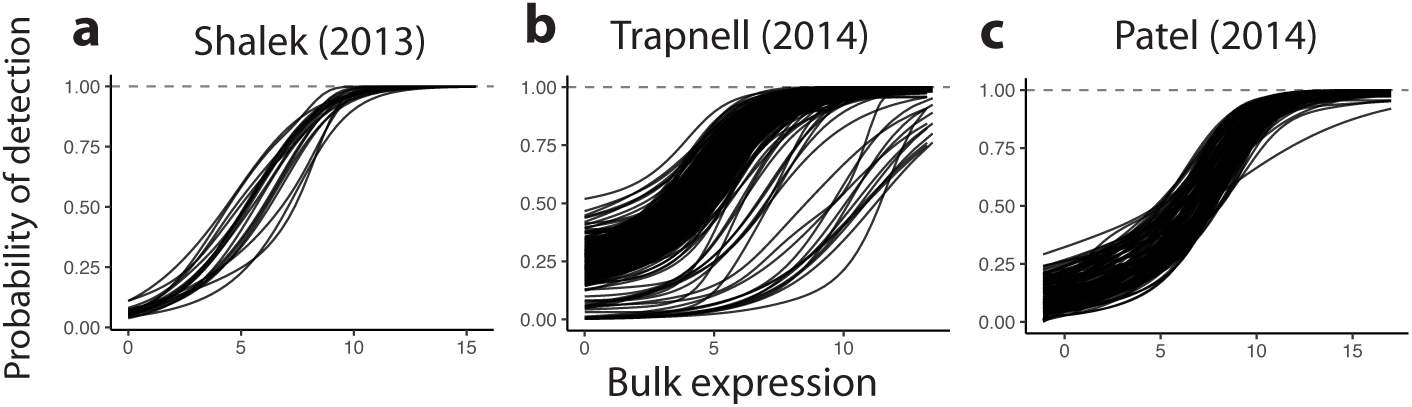
Non-decreasing splines fit to the scatter plot *Z_ng_* versus *log_2_(Mg)* for *M_g_* obtained from: A) Shalek et al. [2013], B) Trapnell et al. [2014], and C) Patel et al. [2014]. Each curve represents a single cell.

Note that if there were no censoring and we set *y*_0_ = 0 and *w_g_* = 0, then letting *σ_y_* → 0 would lead to a model closely related to PCA. If the censoring occurred at random, independently of *η*_*ng*_, then our model would reduce to a variant of Probabilistic Matrix Factorization [Mnih and Salakhutdinov, 2008].

Because we allow a different censoring mechanism for each cell, this model is considerably more complex than previously proposed methods such as ZIFA. To fit the model we leverage the idea of borrowing information across cells to stabilize cell-specific estimates through an empirical Bayesian hierarchical model. We also allow the model to learn the correct latent dimensionality through Automatic Relevance Determination (ARD) [Bishop, 1999]. The Methods section describes the details.

### 2.2 Assessment with published scRNA-Seq data

We used the *two-batches* dataset to demonstrate the improvements provided by our algorithm. In this dataset all cells were extracted from the same biological specimen, with 53 cells processed in one batch and 65 cells processed in another. We applied PCA, t-SNE, and ZIFA with latent dimension *L* = 2, while we initialized *L* = 10 in VAMF. The difference was because *L* is an upper bound on the dimensionality in VAMF but an exact value in other methods. Only the top two VAMF dimensions were used to compare against competing methods.

Since batch should not correlate with new discovered groups, we assessed each method by its ability to avoid predicting batches based on the estimated latent factors. While PCA, t-SNE, and ZIFA all incorrectly detected two clusters (Figure 1), VAMF correctly showed only one cluster (Figure 4A) because it correctly attributed the unwanted variability to differences in detection rates across batches (Figure 4B).

**Figure 4:**
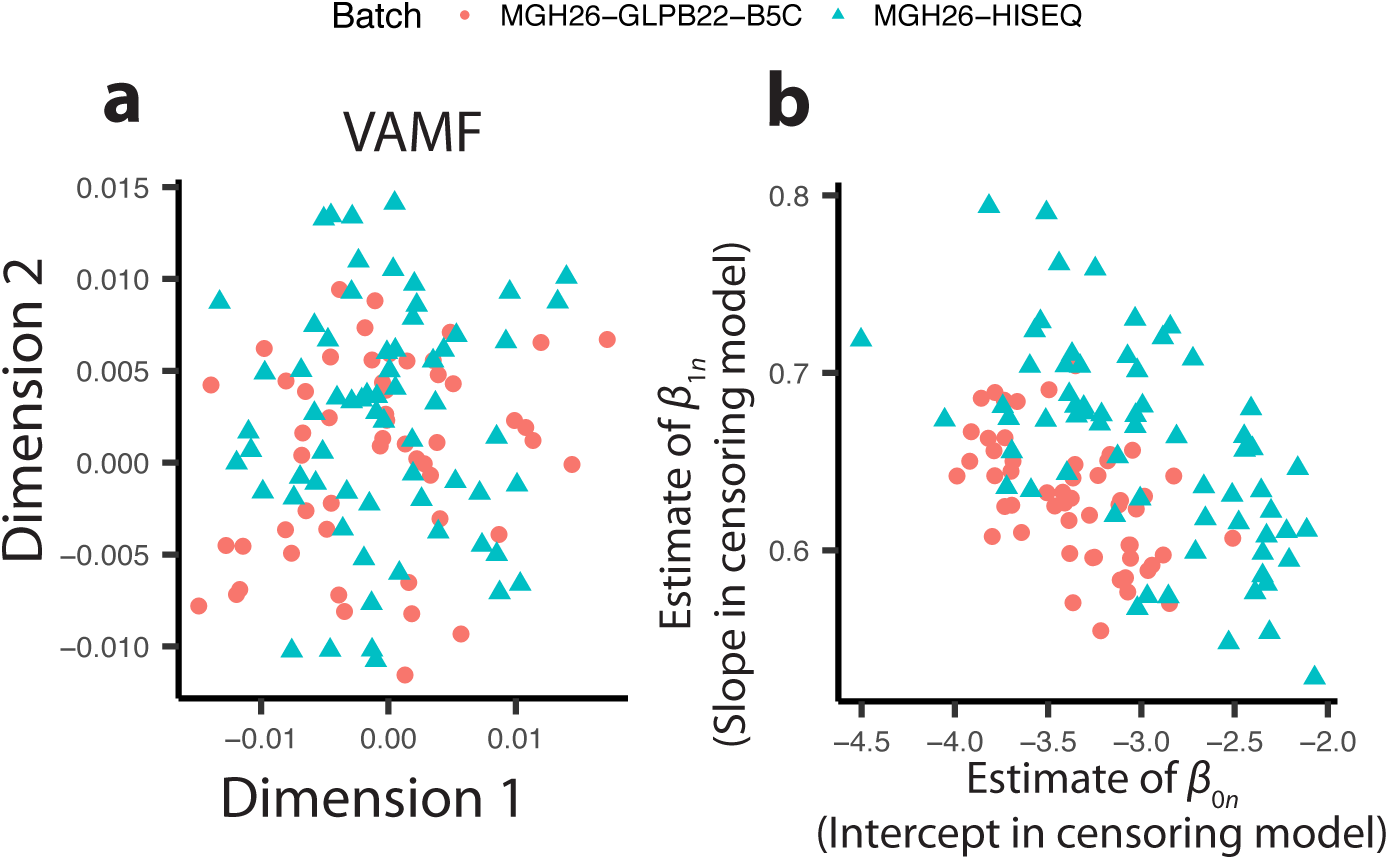
Our method Varying-censoring Aware Matrix Factorization (VAMF) (A) correctly removes differences in the *two-batches* dataset (represented by different shapes and colors) because it correctly attributes the unwanted variability to (B) differences in detection rates across batches, which are captured by the intercept and slope estimates in the censoring model.

To provide a more systematic comparison, we followed Pierson and Yau [2015] and trained linear and quadratic discriminant classifiers (LDA and QDA) on the top two factors estimated by each competing dimension reduction method. We then calculated misclassification error rates as a measure of cluster separability. Because some of the methods we tested were quite computationally intensive if required to analyze matrices with over 20,000 rows, we randomly sub-sampled 100 sets of 2,000 rows (genes). In this assessment, high cluster separability into the groups defined by the sequencer was indicative of a problem since the experimental bias was driving the result. Therefore, the ideal batch separability rate was 0.5: equivalent to guessing. VAMF greatly outperformed all other methods in this assessment (Figure S1), with median batch separability close to 0.65, while other methods had median separability ranging from 0.7 (t-SNE) to 0.9 (PCA and ZIFA). VAMF also outperformed two unpublished methods, Factorial Single Cell Latent Variable Model (f-scLVM Buettner et al., 2016) and Zero-Inflated Negative Binomial Wanted Variation Extraction (ZINB-WAVE Risso et al., 2017), that we included in this comparison.

Additionally, we analyzed the entire *glioblastoma* dataset jointly. We selected cells and filtered for the subset of roughly 6,000 genes that were used in the original analysis by Patel et al. [2014]. Using the same normalization and transformation procedure as described above, we ran all methods to examine whether differences between the batches/tumors changed when adjusting for detection rates using VAMF. VAMF was the only method to remove the batch effect, but did not entirely remove the biological specimen effect (Figure 5).

**Figure 5:**
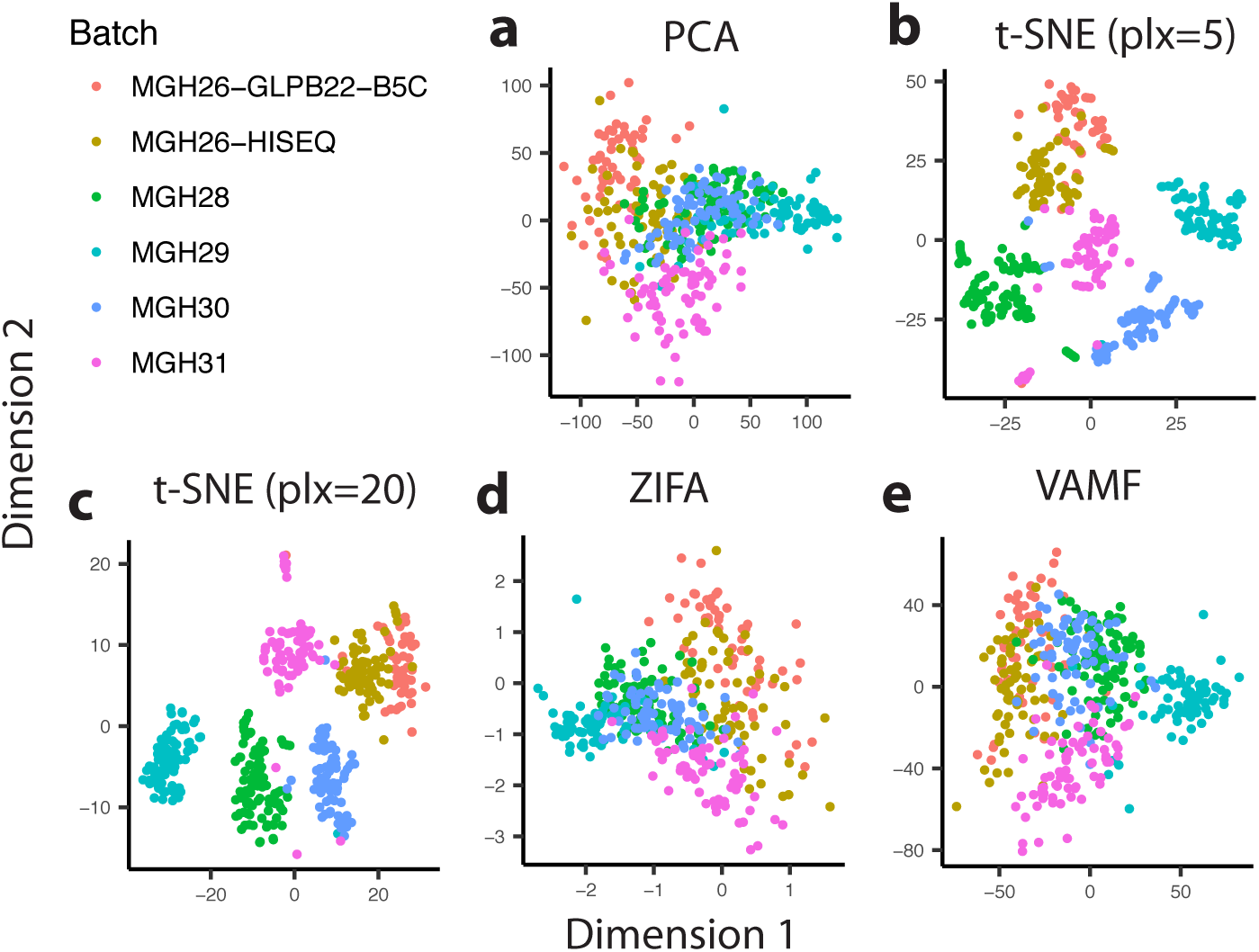
Comparison of dimension reduction methods using all five tumors (about 400 cells) from the *glioblastoma* data. The log of counts from approximately 5800 genes included by the original authors was normalized by SCRAN [L. Lun et al., 2016]. All methods show strong differences between each tumor, but VAMF reduces the distance between the two technical batches of MGH26 to form a single cluster. Log transformation was base two with a pseudocount of one for all methods except VAMF, where no pseudocount was used.

### 2.3 Simulation Study

To further demonstrate that VAMF outperforms other methods, we used simulated data. We generated two scenarios: a *noise only* scenario and a *four biological groups* scenario. In both scenarios, we mimicked batch effects through differences in the detection rates. We compared to what are currently the most widely used methods for dimensionality reduction of scRNA-Seq data, namely PCA, t-SNE and ZIFA.

For the *noise only* scenario, we produced a latent expression matrix *η* with 160 cells and 1,000 genes composed of Gaussian noise with unit standard deviation. Thus, there were no defined biological groups, just random variation. To this we added Gaussian gene effects *w_g_* with standard deviation *σ_w_* = 2. Then, we added a global intercept term of *y*_0_ = 5. Next, we censored the expression values using the logistic model. To mimic a batch effect, we set the values of *β*_0*n*_ so that the cells were divided into a group with high detection rates (*P_n_* ≈ 0.4) and a group with low detection rates (*P_n_* ≈ 0.2). We also randomly set 10% of all data to zero. Such high rates of zeros are commonly observed in experimental scRNA-Seq experiments. The slopes were set to *β*_1*n*_ = 1. In summary, in this scenario all cells had essentially the same expression profiles and differences were artificially introduced through the detection rates, mimicking a common source of technical variation when batch effects are present.

When applied to this simulated data, the varying censoring caused all methods, except VAMF, to recover two clusters (Figure 6A-E). Cells with similar detection rates clustered together. VAMF correctly adjusted for differences in detection rates across cells and showed a latent structure much closer to the actual structure. The detection rate variability was captured by the estimates of *β*_0*n*_ (Figure S2A). VAMF was the only method that avoided the false discovery.

**Figure 6:**
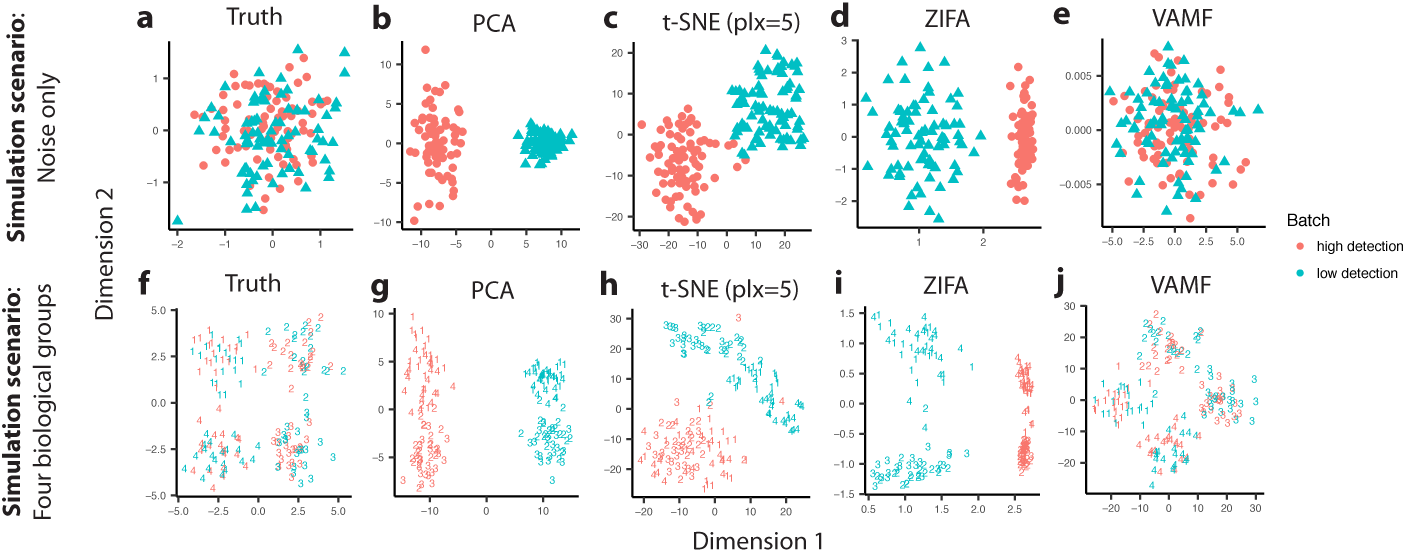
Comparison of dimension reduction methods applied to simulated data: *noise only* scenario (top row) and *four biological groups* scenario (bottom row). Shown are the first two dimensions for (A, F) The true model, (B, G) PCA, (C, H) t-SNE with perplexity (plx) parameter 5, (D, I) ZIFA, and (E, J) our method VAMF. In the *noise only* simulation scenario, a method that does not identify clusters is preferred. High and low detection rate groups are denoted with different colors and symbols. In the *four biological groups* simulation scenario, a method that identifies four clusters corresponding to the four biological groups and does not cluster by the different detection rates is preferred. High and low detection rate groups are denoted with two colors and different latent clusters are represented by numeric symbols. In both simulations, only VAMF accurately recovers the true latent space.

In the second scenario *(four biological groups)*, we introduced variability mimicking differences between undiscovered new cell types. We simulated four clusters of 40 cells each from a mixture of two-dimensional spherical Gaussian distributions, representing the cell type populations. Each cell was represented by its *u_n_* vector of length two. We then projected these into a 50 dimensional space using random loadings *v_g_* to represent the informative, or variable genes. We then concatenated an additional 950 genes with expression values uncorrelated to population classification. To this complete data matrix, we applied the same row, column, and global bias terms as in the *noise only* scenario. We again censored the cells according to the high and low detection groups described above, placing half of the cells from each biological cluster (20 cells) into each of the batches to avoid confounding the technical and biological sources of variation. This scenario mimicked data in which both batch induced technical variability and biological differences represented by different expression profiles were present. Once again, VAMF clearly outperformed all other methods (Figure 6F-J). We repeated this simulation with *G* = 1000 ten times, and also ran ten replicates of a scenario with *G* = 500. For each replicate and scenario, we fit PCA, t-SNE, ZIFA, and VAMF. We then trained four-way LDA classifiers and measured misclassification error. In both scenarios, VAMF had lower error rates than competing methods, indicating successful recovery of the latent clusters despite the varying censoring (Figure S3).

In all scenarios, we specified a dimension of *L* = 2 for t-SNE (with perplexities 5 and 20), PCA, and ZIFA. To illustrate the effect of dimension learning in the VAMF model, we set *L* = 10 and investigated whether VAMF would identify the latent space as being two-dimensional. VAMF learned one-, two-, and three-dimensional spaces in 25%, 60%, and 15% of the latent cluster simulations described above. While exact dimensionality learning is not necessary for identifying latent clusters accurately, VAMF did reasonably well on this much more challenging estimation task.

## 3 Methods

### 3.1 Parameter Estimation

We have posed a model with 2 + *N*(*L* + 2) + *G*(*L* + 1) parameters which we need to fit to *N* × *G* data points, the majority of which are zeros. Unconstrained estimation procedures such as maximum likelihood estimate (MLE) are therefore not appropriate. To impose regularization we applied an empirical Bayesian approach, placing prior distributions on our parameters, and reporting approximate posterior means for the cell factors *u_n_* along with the other variables of interest. Our model parameters were divided into those for which data was highly informative and those for which it was not. For the first group, we used highly uninformative priors to assure the estimates were largely data-driven, similar to MLEs. Specifically we used priors from the Cauchy family with heavy tails as suggested by Gelman [2006]. For the other group of parameters we used Gaussian priors, which enforced stronger regularization. For these, we derived the priors from data.

#### 3.1.1 Latent Model Parameters

We placed a Cauchy prior on the global baseline expression level *y*_0_ centered at the empirical median m of the detected (nonzero) log_2_(*Y_ng_*) values. To ensure a diffuse distribution we used scale parameter 5, which provided essentially a flat prior. For the feature level, effects we used the prior 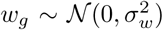 with the standard deviation *σ_w_* derived from the data by computing the median absolute difference (MAD) of the detected genes: MAD(log_2_(*Y_ng_*) − *m*) for *n* such that *Z_ng_* = 1. For the sample factors, we used the prior 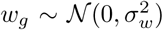. Note that here, the scale hyperparameters for *u_n_* were unnecessary since they could be analytically absorbed into the priors on *v_g_* described next. Automatic relevance determination (ARD) (Bishop, 1999) was used to learn the dimensionality *L* by setting *v_g_* ~ *N*(0, Σ_*v*_) where Σ_*v*_ is a diagonal matrix with elements 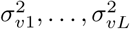 with each *σ_vl_* given a Gamma prior with shape 2 as suggested by Chung et al. 2012]. This family of Gamma distributions has the property that the mode is equal to the scale (inverse rate) parameter. The common scale hyperparameter was determined from the data. Specifically, after centering the data matrix by subtracting off the median as previously described, we computed the scaled MAD values of each row (gene or feature), taking into account only the nonzero elements. Then, the scale hyperparameter was set to the average of these values across all rows. The ARD prior allowed the model to prune away unneeded dimensions during inference by shrinking 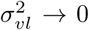. The shape parameter of 2 allowed 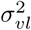 to be arbitrarily close to zero but never exactly zero, preventing numerical singularities [Chung et al., 2012]. A standard half-Cauchy prior was used for the variance of the noise (*σ_y_*).

#### 3.1.2 Censoring Mechanism Parameters

The empirical approach to forming priors for *β*_0*n*_ and *β*_1*n*_ was challenging because although *Z_ng_* was observed, *η_ng_* was not. As described above, bulk RNA-Seq expression measurements *M_g_* provided a useful approximation for *η_ng_* that we used to estimate *β*_0*n*_ and *β*_1*n*_. However, the typical study does not include both single cell and bulk RNA-Seq data from the same specimens because this was done only for assessment purposes during the early testing phase of the single cell technology. However, here we used these assessment experiment data to construct priors for *β*_0*n*_ and *β*_1*n*_ that we then used for future datasets.

To estimate *β*_0*n*_ and *β*_1*n*_ we fit logistic regression to the data from each cell in Shalek et al. [2013], Trapnell et al. [2014], and the *two-batches* dataset [Patel et al., 2014]. We then examined the distributions of the cell-specific MLEs 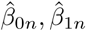 to define reasonable priors for the VAMF model which could be applied to single-cell data lacking a matching bulk RNA-Seq profile.

For the intercept parameters *β*_0*n*_ we leveraged the insight that these parameters are strongly related to detection rate *P_n_*. For example, note that *β*_0*n*_ determines a horizontal shift in the logistic curves and as a consequence, for a fixed value of *β*_1*n*_, the parameter *β*_0*n*_ determines the point *p* at which *f_n_*(*p*) crosses the 0.5 detection probability. We confirmed the strong relationship between these two quantities by comparing each logistic regression MLE 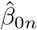 to its corresponding empirical *P_n_* (Figure S2A). This implied that there was in fact information about *β*_0*n*_ in the data and we therefore applied a diffuse Cauchy prior. Furthermore, this motivated an empirical Bayes location hyperparameter for each cell. Specifically, plugging in the prior mean of the slopes *β*_1*n*_ ≈ .5 and a rough estimate of single-cell expression *η*_*ng*_ ≈ *m* into Equation 1 we obtained *P_n_* ≈ *g*(*β*_0*n*_ + .5*m*), where 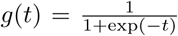 is the inverse logit function. We then used this to obtain a location parameter for the Cauchy prior of *g*^−1^ (*P_n_*) − .5*m*. We set the scale to 1 based on examining the distribution of MLEs 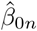 described previously. Our choice of location parameters encouraged the model to use intercepts that were consistent with the observed detection rates in the data at hand. However, by using a Cauchy prior we permitted the model to accommodate the unexpected and highly variable detection rates seen in practice.

Slope parameters *β*_1*n*_ appeared to be normally distributed with a mean of about 0.5 and a standard deviation of about .1 (Figure S2B). Hence we used this distribution as an informative prior in the VAMF model.

We fit our hierarchical model with the Automatic Differentiation Variational Inference (ADVI) algorithm [Kucukelbir et al., 2015] implemented in Stan [Carpenter et al., 2016]. We refer to this method for estimating factors in the presence of censored data as Variable-censoring Aware Matrix Factorization (VAMF).

### 3.2 Rotations of Factors for Improved Interpretability

The joint likelihood model described above yields cell and gene factors *u_n_, v_g_* that are combined by an inner product, such that the likelihood is invariant to rotations of the factors. Furthermore, there is no guarantee the factors are not correlated with each other, and factors are not ordered by importance. To alleviate this and facilitate interpretability, such as comparisons with PCA, we orthogonalize and rotate the factors by the following procedure.

Specifically, we let *U* be the *L* × *N* matrix formed by stacking the posterior means of *u_n_* cell factors into the columns. Let *V* be the *L* × *G* matrix of stacked posterior means of gene factors *v_g_*. The rows of *V* form a basis for the *L* ≪ *G* dimensional subspace learned by VAMF in the context of the *G* dimensional original data space. Furthermore, the likelihood depends on *U* and *V* only through *V′U*. Hence, let *V* = *ADQ′* be the singular value decomposition where *A* and *D* are *L* × *L* and *Q* is *G* × *L*. Then the likelihood depends on *QŨ* = *Q*(*DA′U*). Now, the columns of *Q* are orthonormal and span the same space as the rows of *V*. The vectors *ũ_n_* = *DA′u_n_* are in the columns of *Ũ* and represent rotated and scaled versions of the original *u_n_* factors. After this transformation, the rows of *Ũ* are analogous to principal components and are in decreasing order according to the *L*_2_ norm (assuming the standard decreasing ordering of singular values in *D*). We can inspect these *L*_2_ norms to determine the inferred dimension of the subspace; components with large norms are dominant axes of variation, whereas components with small norms are extraneous dimensions that have been pruned away by ARD.

## 4 Discussion

We have demonstrated the effectiveness of Varying-censoring Aware Matrix Factorization (VAMF) for performing dimension reduction in the presence of cell-specific varying detection rates on simulated and real datasets. While differences in detection rates are often the result of differing technical experimental conditions, in other cases they may be due to real biology. For example, in studies of embryological development one may expect the number of detected genes to increase over time [Deng et al., 2014], it provides users with a decomposition of variability into that arising from the detected genes and that arising from the differences in detection rates. Specifically, after applying our VAMF model, users can visualize or apply clustering algorithms to the cell factors *u_n_* to discover new cell types with differing expression profiles and they can visualize the posterior estimates of *β*_0*n*_ to study the severity of censoring across cells. We also note that if batch effects alter the detected measurements through mechanisms other than detection rates, our model will capture these as *u_n_*. It is up to the user to determine if further adjustments are needed. For example, in Figure 5E no dimension reduction method can clarify whether the clustering by biological group may in fact be an unwanted batch effect since the experiment confounded sequencing batches with biological groups.

While the VAMF method was the most accurate in our assessments, it was computationally expensive. Like other informative censoring models such as ZIFA, the algorithm implicitly requires imputing the missing values in the entire data matrix at each iteration. In contrast, models assuming zeros occur at random may compute over only the observed data values resulting in faster algorithms. Models like t-SNE that use only pairwise distance metrics between points also save on computation. However, as we have shown these will lead to potentially false discoveries.

Future work should focus on scaling VAMF to larger datasets to improve speed and memory usage. We also found some advantages from some of the unpublished methods we compared to. Although these methods failed to capture cell-to-cell detection rate differences, they have added features that could be incorporated into VAMF. For example, the VAMF censoring mechanism is a generalization of the “Hurdle Model” used by the unpublished f-scLVM method [Buettner et al., 2016]. The f-scLVM model has the advantage of using gene set annotations to learn latent factors, and separates biological and technical factors by assuming the former are sparse while the latter are dense. It should be possible to combine the VAMF censoring mechanism with the f-scLVM factor model, which could lead to an improvement in performance for both methods.

## 5 Software

The code used to produce results for this paper, including an implementation of VAMF, is available at: https://github.com/willtownes/vamf-paper.

## Acknowledgments

The authors thank Cem Sievers, Keegan Korthauer, Jeff Miller, and Mike Love for valuable feedback. *Conflict of Interest:* None declared.

## A Supplementary Methods

### A.1 Bioinformatics Preprocessing

Metadata were obtained from Gene Expression Omnibus (GEO; Edgar et al., 2002) for the three datasets Shalek et al. [2013], Trapnell et al. [2014], Patel et al. [2014]. For the first two of these, we simply used the expression matrices posted to GEO by the original authors, which had already been normalized to TPM and FPKM respectively. For the data of Patel et al. [2014], we could not use the authors’ normalization scheme since it erased the zeros and did not include all genes nor all cells. We therefore started from FASTQ files downloaded from Sequence Read Archive (SRA; https://www.ncbi.nlm.nih.gov/sra). We obtained estimated counts for ENSEMBL genes using Kallisto [Bray et al., 2015].Full details and count tables, including all genes and all samples, are available at: https://github.com/willtownes/patel2014gliohuman.

### A.2 Computational Performance and Software Versions

While PCA and t-SNE were unmatched for speed, they do not address censoring. Our method VAMF had computational performance comparable to ZIFA on smaller datasets and superior performance on a larger dataset (Table S1) even though we specified a larger number of latent dimensions, which would be expected to slow down the inference procedure. Analyses were run on a Mac Mini with 2.6 GHz Intel Core i5 processor and 16 Gb of memory and MacOS Sierra operating system. The R version was 3.4.0.

**Table S1:**
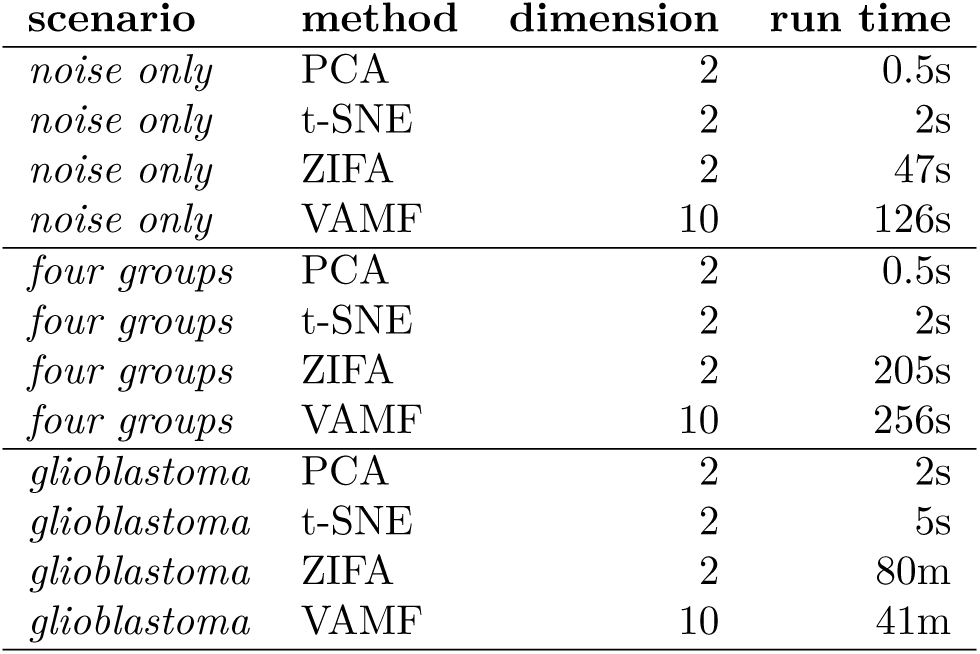
Comparison of computational performance of PCA, t-SNE, ZIFA, and VAMF methods on the *noise only* simulation scenario, the *four biological group* simulation scenerio, and the real *glioblastoma* data set. ‘s’ seconds, ‘m’ minutes

| scenario | method | dimension | run time |
| --- | --- | --- | --- |
| *noise only* | PCA | 2 | 0.5s |
| *noise only* | t-SNE | 2 | 2s |
| *noise only* | ZIFA | 2 | 47s |
| *noise only* | VAMF | 10 | 126s |
| *four groups* | PCA | 2 | 0.5s |
| *four groups* | t-SNE | 2 | 2s |
| *four groups* | ZIFA | 2 | 205s |
| *four groups* | VAMF | 10 | 256s |
| *glioblastoma* | PCA | 2 | 2s |
| *glioblastoma* | t-SNE | 2 | 5s |
| *glioblastoma* | ZIFA | 2 | 80m |
| *glioblastoma* | VAMF | 10 | 41m |

R package **scam** was used to fit monotone splines. For all methods, rows and columns composed of entirely zeros were removed from the data before processing. For PCA and t-SNE, data were centered and scaled prior to analysis. We used R function **prcomp** for PCA and R package **Rtsne** for t-SNE [Krijthe, 2017].

## Supplementary Figures and Tables

**Figure S1:**
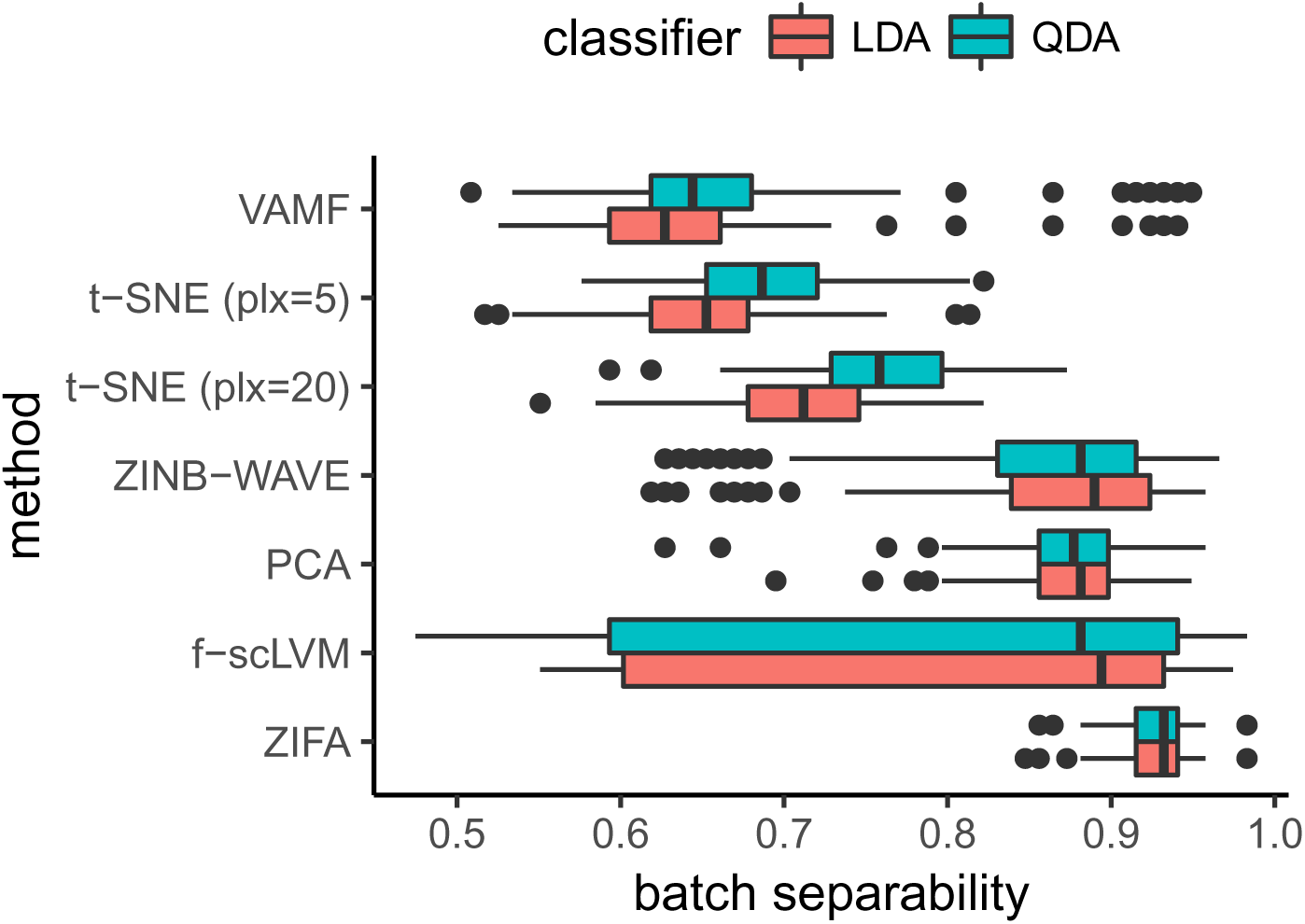
VAMF is more effective at removing technical batch effects mediated through detection rates than competing methods using the *two-batches* dataset. Using 100 replicates of 2,000 random genes in the Patel et al. [2014] tumor MGH26 (118 cells in two technical batches), we applied each dimension reduction method and then trained linear (LDA) and quadratic (QDA) discriminant analysis classifiers on the first two latent dimensions using the known batch labels. Batch separability was quantified using one minus the misclassification error rate. High separability implies inability to remove batch effects in this dataset. We show results for VAMF, t-SNE with perplexity (plx) parameters 5 and 20, Zero-Inflated Negative Binomial Wanted Variable Extraction (ZINB-WAVE), PCA, Factorial Single Cell Latent Variable Model (f-scLVM), and ZIFA.

**Figure S2:**
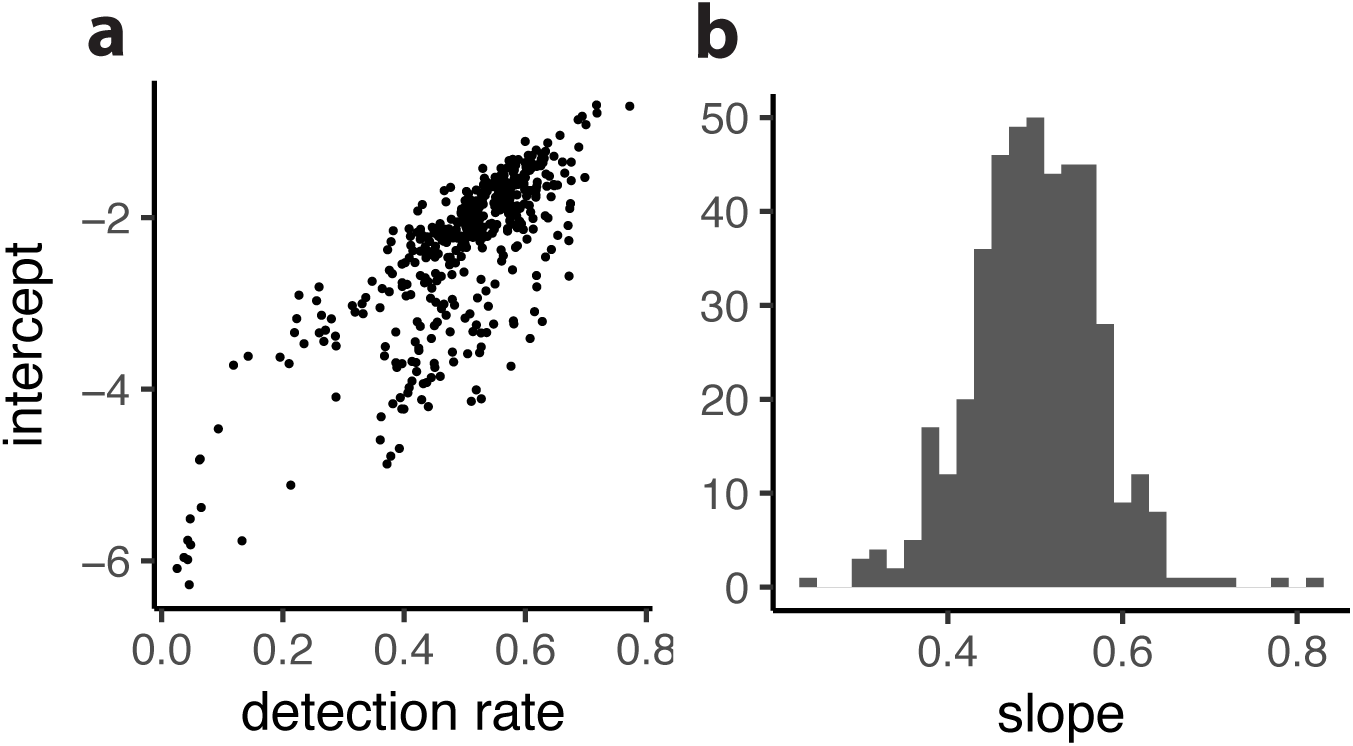
We implemented an empirical approach to define the priors for the intercept and slope parameters, *β*_0*n*_ and *β*_1*n*_, for the censoring mechanism logistic curves. We obtained maximum likelihood estimates (MLEs) for these parameters using the three assessment datasets. A) MLEs for the intercept parameters *β*_0*n*_ plotted against the observed detection rate show relatively strong correlation. We used this to define a prior expected value for β_0n_. B) The MLEs for the slope parameters *β*_1*n*_ appear to be approximately normally distributed with expected value 0.5 and standard deviation 0.1. We used this as the prior distribution for the slope parameters.

**Figure S3:**
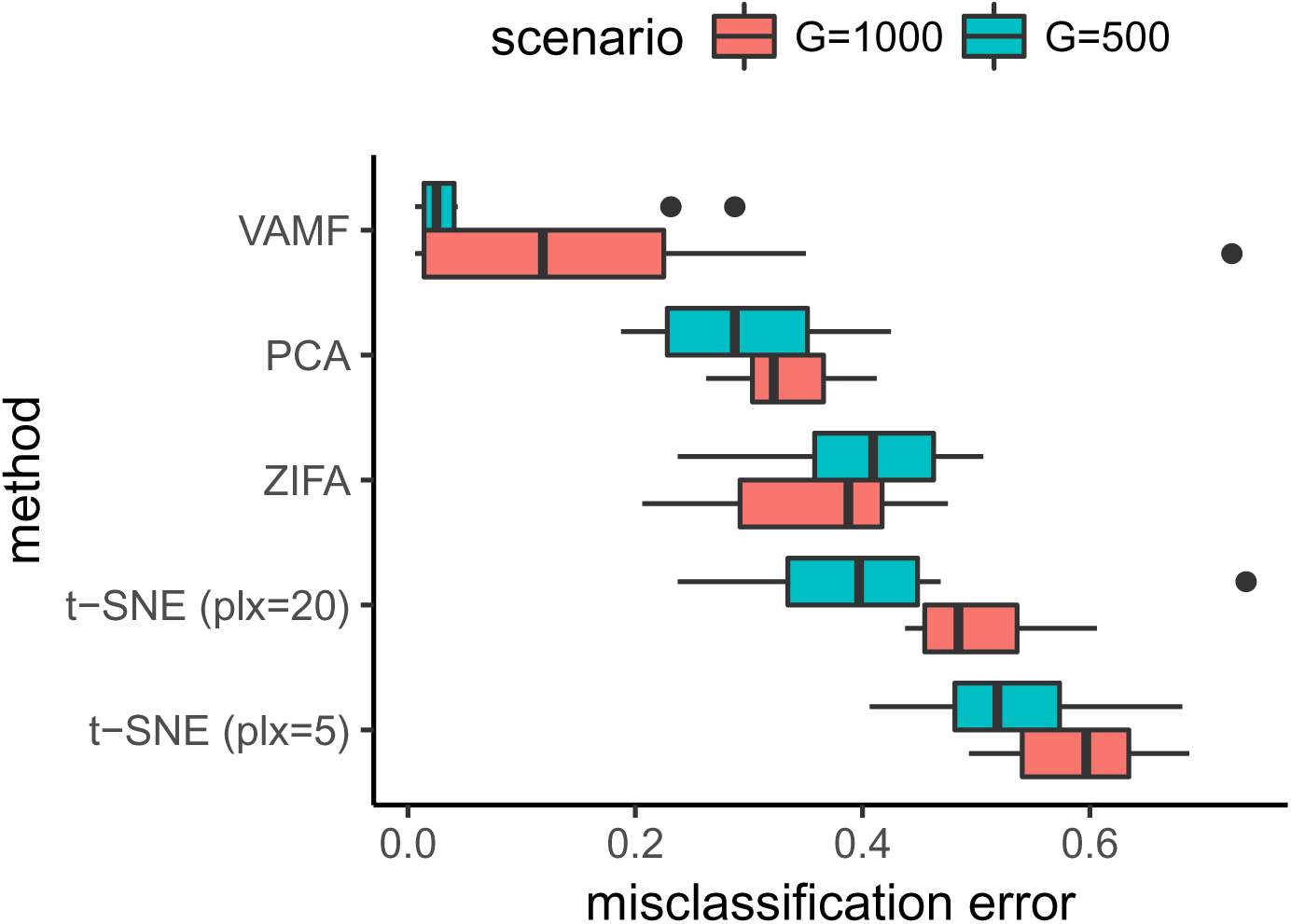
VAMF is more effective at recovering latent clusters than competing methods. We simulated ten replicates each of two scenarios *G* = 500 and *G* = 1000 total genes with 50 informative genes. After obtaining latent factors from VAMF, PCA, ZIFA, and t-SNE with perplexity (plx) parameters 5 and 20, we trained four-way Linear Discriminant Analysis classifiers and calculated misclassification error rates using ground truth labels and the top two latent dimensions. A low error rate indicates better accuracy in recovery of the latent clusters.

